# Transposons contribute to the functional diversification of the head, gut, and ovary transcriptomes across *Drosophila* natural strains

**DOI:** 10.1101/2022.12.02.518890

**Authors:** Marta Coronado-Zamora, Josefa González

**Affiliations:** Institute of Evolutionary Biology, CSIC, UPF

## Abstract

Transcriptomes are dynamic, with cells, tissues, and body parts expressing particular sets of transcripts. Transposons are a known source of transcriptome diversity, however studies often focus on a particular type of chimeric transcript, analyze single body parts or cell types, or are based on incomplete transposon annotations from a single reference genome. In this work, we have implemented a method based on *de novo* transcriptome assembly that minimizes the potential sources of errors while identifying a comprehensive set of gene-TE chimeras. We applied this method to head, gut and ovary dissected from five *Drosophila melanogaster* natural strains, with individual reference genomes available. We found that ∼19% of body part-specific transcripts are gene-TE chimeras. Overall, chimeric transcripts contribute a mean of 43% to the total gene expression, and they provide DNA binding, catalytic activity, and DNA polymerase activity protein domains. Our comprehensive dataset is a rich resource for follow-up analysis. Moreover, because transposable elements are present in virtually all species sequenced to date, their relevant role in spatially restricted transcript expression is likely not exclusive to the species analyzed in this work.

## INTRODUCTION

In contrast to the genome, an animal’s transcriptome is dynamic, with cell types, tissues and body parts expressing particular sets of transcripts (Pan et al. 2008; Barash et al. 2010; Brown et al. 2014; Söllner et al. 2017). The complexity and diversity of the transcriptome arises from the combinatorial usage of alternative promoters, exons and introns, and polyadenylation sites. A single gene can, therefore, encode a rich repertoire of transcripts that can be involved in diverse biological functions, and contribute to adaptive evolution and disease (*e.g.*, Kiyose et al. 2022; Verta and Jacobs 2022; Marasca et al. 2020; Singh and Ahi 2022). The potential contribution of transposable element (TE) insertions to the diversification of the transcriptome was analyzed soon after the first whole-genome sequences were available (Franchini et al. 2004; Ganko et al. 2003; Jordan et al. 2003; Lipatov et al. 2005; van de Lagemaat et al. 2003). TEs are present in virtually all genomes studied to date, are able to insert copies of themselves in the genome and, although their mutation capacity is often harmful, they also represent an important source of adaptive genetic variation (Casacuberta and González 2013; Cowley and Oakey 2013; Schrader and Schmitz 2019; Volff 2006). While TEs are a known source of transcriptome diversity, the majority of studies so far rely on incomplete transposon annotations from a single reference genome (*e.g.*, Lipatov et al. 2005). Moreover, methodologies are often specifically designed for particular types of chimeric gene-TE transcripts, *e.g.*, TE-initiated transcripts (Babaian et al. 2019), particular types of TEs, *e.g.,* L1 chimeric transcripts (Pinson et al. 2018), or have been applied to individual cell types or body parts, (*e.g.*, Babarinde et al. 2021; Treiber and Waddell 2020). As such, our knowledge on the contribution of TEs to transcriptome diversification is still partial.

Two of the most studied mechanisms by which TEs can generate chimeric transcripts are by providing alternative promoters and protein domains. In human and mouse, 2.8% and 5.2% of the total transcript start sites occur within retrotransposons, respectively (Faulkner et al. 2009). In *D. melanogaster*, over 40% of all genes are expressed from two or more promoters, with at least 1,300 promoters contained in TEs (Batut et al. 2013). As well as individual examples of TEs providing protein domains (Cordaux et al. 2006; Newman et al. 2008; Tipney et al. 2004), a comparative genomic analysis of tetrapod genomes revealed that capture of transposase domains is a recurrent mechanism for novel gene formation (Cosby et al. 2021). There is also evidence for the retrotransposon contribution to protein novelty. Approximately 9.7% of endogenous retrovirus open reading frames across 19 mammalian genomes evolve under purifying selection and are transcribed, suggesting that they could have been co-opted as genes (Ueda et al. 2020). Across insects, and depending on the methodology used, the percentage of newly emerged domains (<225 mya) due to TEs was estimated to be from 1.7% to 6.6% (Klasberg et al. 2018). However, studies that identify and characterize a comprehensive set of gene-TE chimeras to provide a complete overview of their contribution to both transcriptome and protein diversification are still missing.

Besides describing the diverse contributions of TEs to the transcriptome, analyzing the relative contribution of gene-TE chimeras to the total gene expression is highly relevant, as it is informative of the potential functional relevance of the transcripts identified. Studies performed so far suggest that this contribution is related to the position of the TE in the transcript. Transcripts with a TE inserted in the 5’UTR or internal coding exons show significantly lower mean levels of expression compared with non-chimeric TE-gene transcripts (Babarinde et al. 2021). TEs inserted in 3’UTRs were associated with reduced gene expression both in humans and mice, but with increased gene expression in human pluripotent stem cells (Babarinde et al. 2021; Faulkner et al. 2009). In addition, whether specific TE types contribute to tissue-specific expression has been explored in mammals, where retrotransposons were found to be overrepresented in human embryonic tissues (Conley et al. 2008; Faulkner et al. 2009). In *D. melanogaster*, the contribution of TEs to tissue-specific expression has only been assessed in the head, with 833 gene-TE chimeric genes described (Treiber and Waddell 2020). Thus, whether the contribution of chimeric gene-TE transcripts is more relevant in the *D. melanogaster* head compared with other body parts is still an open question.

Within genes, TEs could also affect expression by changing the epigenetic status of their surrounding regions. In Drosophila, repressive histone marks enriched at TEs spread beyond TE sequences, which is often associated with gene down-regulation (Lee and Karpen 2017). However, there is also evidence that TEs containing active chromatin marks can lead to nearby gene overexpression (Guio et al. 2018). Genome-wide, the joint assessment of the presence of repressive and active chromatin marks has been restricted so far to the analysis of four TE families (Rebollo et al. 2012) and has never been carried out in the context of chimeric gene-TE transcripts.

In this work, we performed a high-throughput analysis to detect, characterize, and quantify chimeric gene-TE transcripts in RNA-seq samples from head, gut, and ovary dissected from the same individuals belonging to five natural strains of *D. melanogaster* (Figure 1A; Rech et al. 2022). We implemented a method based on *de novo* transcriptome assembly that (i) minimizes the potential sources of errors when detecting chimeric gene-TE transcripts; and (ii) allows to identify a comprehensive dataset of transcripts rather than focusing on particular types (Figure 1B; Lanciano and Cristofari 2020). Additionally, we assessed the coding potential and the contribution of chimeric transcripts to protein domains and gene expression as proxies for their integrity and functional relevance. Finally, we took advantage of the availability of ChIP-seq data for an active and a repressive histone mark, H3K9me3 and H3K27ac, respectively obtained from the same biological samples to investigate whether the TEs that are incorporated into the transcript sequences also affect their epigenetic status.

**Figure 1.**
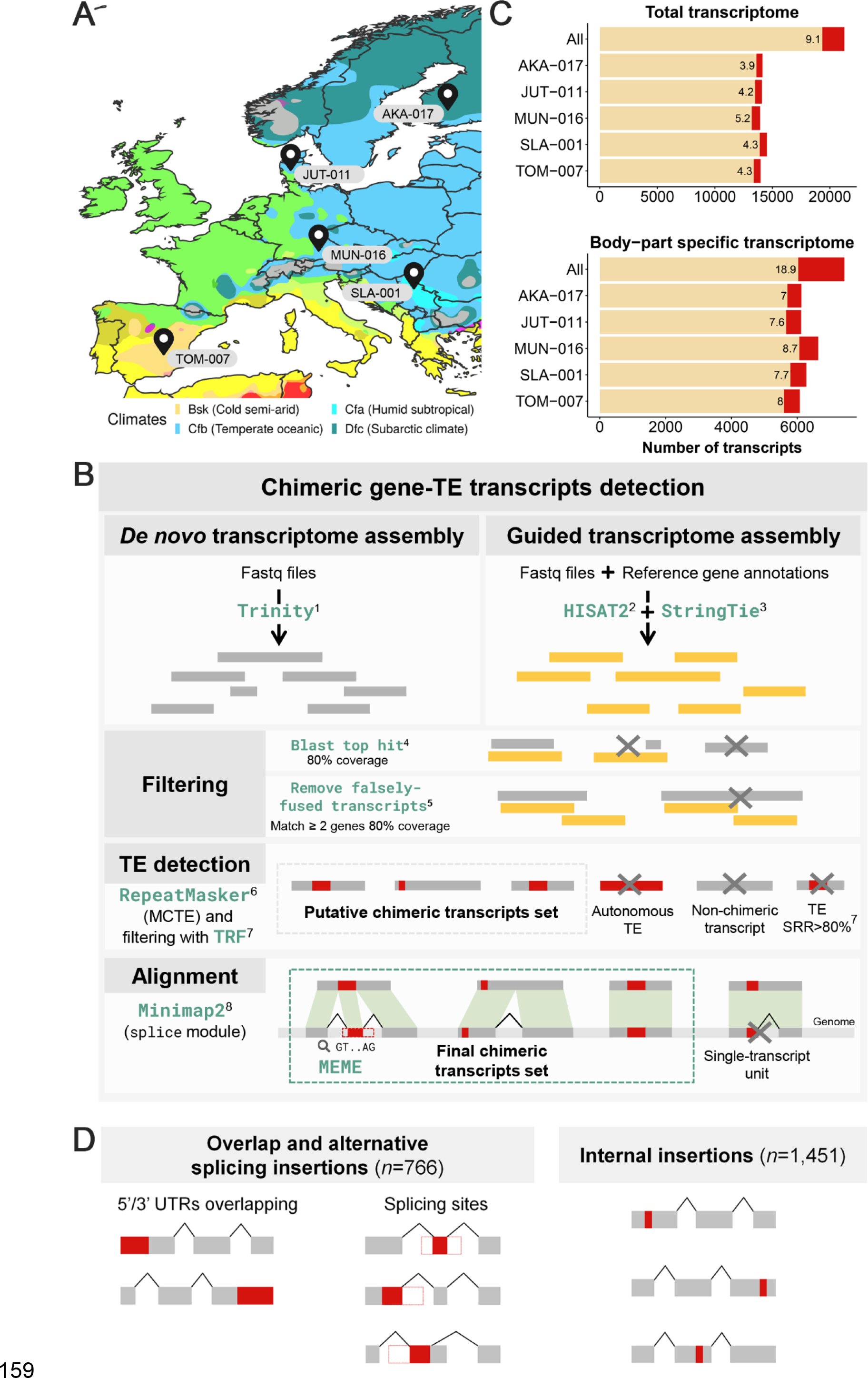
Detection of chimeric gene-TE transcripts in five strains of *D. melanogaster*.**A.** Map showing the sampling locations of the five European strains of *D. melanogaster* used in this study. TOM-007: Tomelloso, Spain (*BSk*); MUN-016: Munich, Germany (*Cfb*); JUT-011: Jutland, Denmark (*Cfb*); SLA-001: Slankamen, Serbia (*Cfa*); and AKA-017: Akaa, Finland (*Dfc*). Colors represent the climate zones according to the Köppen-Geiger climate distribution (Peel et al. 2007). **B.** Pipeline to detect chimeric transcripts. Two types of transcriptome assembly were performed: a *de novo* assembly using Trinity^1^ (Grabherr et al. 2011) and a genome-guided transcriptome assembly using HISAT2^2^ (Kim et al. 2019) and StringTie^3^ (Pertea et al. 2015). Only *de novo* transcripts that had a minimum 80% coverage with a known transcript were considered to be screened for TEs insertions^4^. Falsely-fusioned transcripts derived from gene-dense compact genomes were removed^5^. RepeatMasker^6^ (Smit et al. 2013) is used with a manually curated TE library (Rech et al. 2022) to detect TEs in the *de novo* assembled transcripts. Transcripts with TE fragments having simple repeats in over 80% of their length were excluded^7^ (Benson 1999). An alignment against the reference genome of each strain is used to define the exon-intron boundaries of transcripts and to identify the position of the TE in the transcript (Li 2018)^8^. Transcripts fully annotated as a TE or detected as single-transcript units when the sequence matched a multi-exonic transcript are discarded. **C.** Contribution of chimeric gene-TE transcripts to the total transcriptome and the body part- specific transcriptome globally and by strain. *All* includes all the transcripts assembled in the three body parts and the five strains. **D.** Schematic of the two groups of chimeric transcripts identified. *Overlap and alternative splicing (AS) insertions* group, and *internal insertions* group. Note that these numbers total more than 1,931 because some chimeric transcripts can have different insertions in different samples. Gray boxes represent exons, red boxes represent a TE fragment incorporated in the mRNA, white boxes represent a TE fragment that is not incorporated in the final mRNA. The black lines connecting the exons represent the splicing events.

## RESULTS

### 9% of D. melanogaster transcripts, across body parts and strains, are gene-TE chimeras

We performed a high-throughput analysis to detect and quantify chimeric gene-TE transcripts in RNA-seq samples from head, gut, and ovary, in five *D. melanogaster* strains collected from natural populations (Figure 1A). The three body parts were dissected from the same individuals, and an average coverage of 32x (22x to 43x) per RNA-seq sample was obtained (3 replicates per body part and strain, Supplemental Table S1; Coronado-Zamora et al. 2023). We *de novo* assembled transcripts in which we annotated TE insertions using a new *D. melanogaster* manually curated TE library (Rech et al. 2022). We only considered *de novo* transcripts that overlap with a known transcript obtained from a reference-guided assembly (Figure 1B), and we removed falsely-fusioned transcripts applying the algorithm described in Lima et al. (2017). We also excluded chimeric transcripts with TE fragments whose sequence contains simple repeats in more than >80% of their length (see *Methods*; Benson 1999; Rech et al. 2022). We then used the reference genome of each strain to define the exon-intron boundaries of each transcript and to identify the position of the TE in the transcript (Figure 1B). The alignment with the reference genome and the accurate TE annotation also allowed us to discard single-unit transcripts, indicative of pervasive transcription when the sequence matched a multi-exonic transcript, and TE autonomous expression, which are two important sources of errors when quantifying the contribution of TEs to gene novelty (Figure 1B; Lanciano and Cristofari 2020).

Overall, considering all the transcripts assembled in the three body parts and the five strains, we identified 1,931 chimeric gene-TE transcripts belonging to 826 genes (Supplemental Table S2A). Thus, approximately 9% (1,931/21,270) of *D. melanogaster* transcripts contain exonic sequences of TE origin. In individual strains, this percentage ranged between 3.9% to 5.2% (549-725 chimeric transcripts per genome) indicating that most of the chimeric gene-TE transcripts are strain-specific (1,312/1,931, 68%), as expected given that the majority of TEs are present at low population frequencies (Figure 1C; Rech et al. 2022). Almost half of the strain-specific chimeric transcripts (48%, 634/1,312) were generated through strain-specific TE insertions, while the other 52% through polymorphic (141) or fixed (599) insertions, indicating that differential exon usage is an important mechanism in generating chimeric transcripts. While the overall contribution of TEs to the transcriptome is 9%, TEs contribute approximately 19% (1,406/7,435) of the total amount of body part-specific transcripts (Figure 1C).

We identified two groups of chimeric gene-TE transcripts (Figure 1D). The first group contains chimeric transcripts which have a TE overlapping with the 5’UTR, the 3’UTR, or introducing alternative splice (AS) sites (*overlap and AS insertions* group: 766 chimeric transcripts from 415 genes). We found that the majority of the TEs in the *overlap and AS insertions* group were adding canonical splice sites motifs (84.1%: 132/157) (Supplemental Table S2B). The second group contains chimeric gene-TE transcripts in which the TE is annotated completely inside the UTRs or internal exons (*internal insertions* group: 1,451 transcripts from 638 genes) (Figure 1D). We hypothesized that this group could be the result of older insertions that have been completely incorporated into the transcripts. Indeed, we found that TEs in this group are shorter than those of the *overlap and AS insertion* group, as expected if the former are older insertions (78% *vs*. 22.7% proportion of chimerics with TEs < 120 bp; χ² test, *p*-value < 0.001; Supplemental Fig. S1; see *Methods*). Additionally, while the majority of gene-TE transcripts in the *overlap and AS insertions* group were strain-specific, we found more transcripts shared between strains than strain-specific in the *internal insertions* group (χ² test, *p*-value < 0.001; Supplemental Fig. S2A and Supplemental Table S2C). This observation is also consistent with this group being enriched for older insertions, and remained valid when we removed the shorter insertions (χ² test, *p*-value < 0.001; Supplemental Table S2C).

To test whether the *overlap and AS insertions* and the *internal insertions* groups contribute differently to the diversification of the transcriptome, we performed all the subsequent analyses considering all the chimeric transcripts together, and the two groups separately. In addition,because shorter insertions might be enriched for false positives, *i.e.,* not corresponding to real TE sequences due to the difficulty of annotating these repetitive regions, we also performed the analysis with the subset of chimeric gene-TE transcripts that contains a fragment of a TE insertion that is ≥120bp (638/766 and 672/1,451 for the *overlap and AS insertions* and the *internal insertions* groups, respectively; see *Methods*).

### Gene-TE chimeric transcripts are more abundant in the head and ovary

Using high-throughput methodologies, 833 chimeric genes were identified in the *D. melanogaster* head (Treiber and Waddell 2020), however, the relative amount of chimeric gene-TE transcripts across body parts has never been assessed before. We found that the majority of the assembled chimeric gene-TE transcripts across the five strains analyzed were body part-specific (72.8%: 1,406/1,931), with only 9.3% (180) shared across all three body parts (Figure 2A and Supplemental Table S3A). The same pattern was found for the *overlap and AS insertions* group and for the *internal insertions* group, when considering all insertions and those ≥120bp (Supplemental Fig. S2B and Supplemental Table S3A).

**Figure 2.**
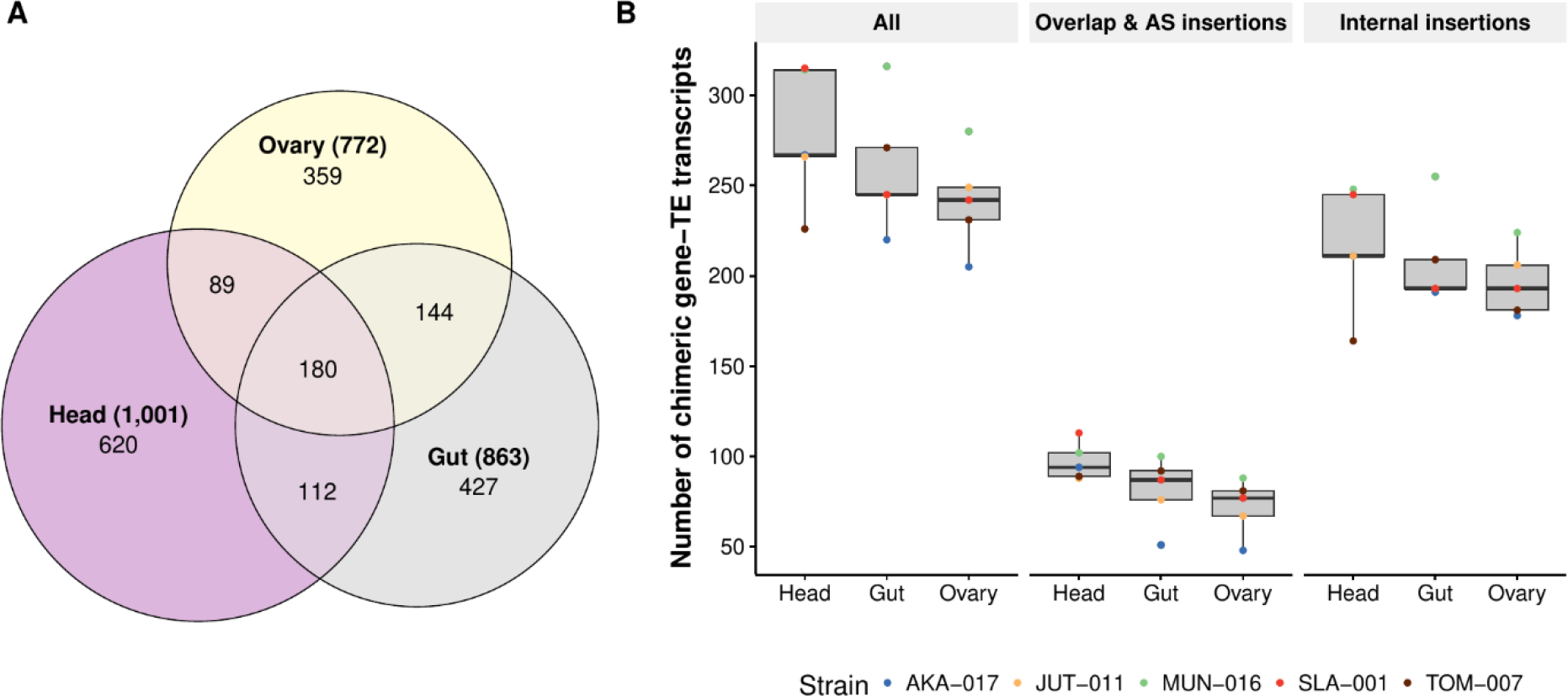
Distribution of chimeric transcripts across body parts and insertion groups. **A.** Venn diagram showing the overall number of chimeric transcripts shared across body parts in the five strains. **B.** Number of chimeric gene-TE transcripts detected by body part, strain and insertion group. *All* includes all chimeric transcripts detected in all body parts and strains.

Head was the body part expressing the most chimeric transcripts (1,001) followed by gut (863) and ovary (772) (Figure 2A and Supplemental Table S3A). Note that 145 of the chimeric genes identified in this work were previously described by Treiber and Waddell (2020), an expected low number given that many of the detected gene-TE chimeras are strain-specific and that the applied methodologies are different. After accounting for differences in the total number of transcripts assembled in each body part, we observed that the head was expressing the same proportion of gene-TE chimeras as the ovary (6.2% head *vs*. 6.2% ovary; χ² test, *p*-value = 1); while head and ovary express more than the gut (5.6% gut; χ² test, *p*-values = 0.032 and 0.048, respectively; Supplemental Table S3C). A similar proportion of chimeric transcripts in head and ovary was also found when the *overlap and AS insertions* and the *internal insertions* groups were analyzed separately, also if we focus on ≥120bp insertions (Figure 2B and Supplemental Table S3C). Overall, the same pattern was also found at the strain level, except for some comparisons (Supplemental Table S3C).

Finally, the head was the body part that expressed the most body part-specific chimeric transcripts (62% head *vs.* 50% gut; χ² test, *p*-value = 0.005, and *vs.* 46.5% ovary, *p*-value < 0.001), while no differences were found between gut and ovary (46.5% ovary *vs.* 50% gut; χ² test, *p*-value = 0.8; Figure 2A). In the three body parts, these proportions were higher than the total proportion of body part-specific transcripts (23.2%, 15.14% and 11.25%, for head, gut and ovary respectively; χ² test, *p*-values < 0.001 for all comparisons; Supplemental Table S3B).

### Most chimeric transcripts contain TE insertions in the 3’UTRs

Chimeric gene-TE transcripts are enriched for TE insertions located in the 3’UTRs in *D. melanogaster* and in mammals (Babarinde et al. 2021; Lipatov et al. 2005; van de Lagemaat et al. 2003). Consistently, we also found that most of the chimeric gene-TE transcripts contain a TE in the 3’UTR (963 transcripts from 417 genes) followed by internal exons (569 transcripts from 266 genes) and insertions in the 5’UTRs (474 transcripts from 273 genes). We also detected 125 chimeric transcripts belonging to 95 genes consisting of a single-unit transcript. Note that 21 of the 5’UTR insertions detected in this work were experimentally validated in a previous analysis that estimated the promoter TE usage across developmental stages in *D. melanogaster* (Batut et al. 2013). Indeed, the number of chimeric genes with a TE inserted in the 3’ and 5’UTRs is higher than expected when taking into account the proportion of the genome that is annotated as UTRs, while there is a depletion of TEs in internal exons (χ² test, *p*-value < 0.001 in the three comparisons; Supplemental Table S4A). It has been hypothesized that the higher number of insertions in 3’UTRs could be explained by lack of selection against insertions in this gene compartment (Jordan et al. 2003; Lipatov et al. 2005). We thus tested whether 3’UTR chimeric transcripts were enriched for TE insertions present in more than one genome, but we found that this was not the case (χ² test, *p*-value = 0.9731; Figure 3A and Supplemental Table S4B).

**Figure 3.**
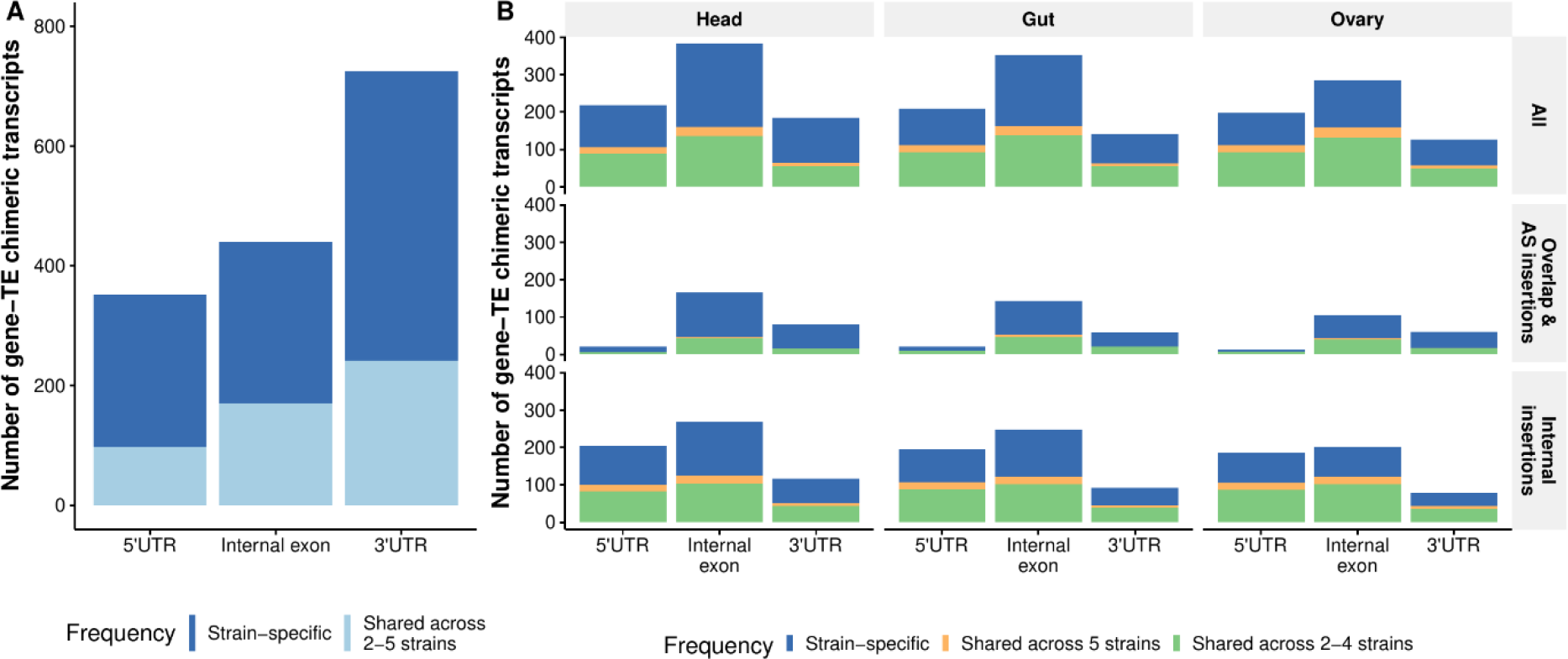
Position and frequency distribution of TEs in chimeric transcripts. A. Number of gene-TE chimeric transcripts by position and frequency**. B.** Number of chimeric gene-TE transcripts by insertion group and body part, according to the insertion position (5’/3’UTRs or internal exons) and frequency. Each dot represents the number of chimeric gene-TE transcripts according to the frequency: strain-specific (blue), shared across two to four strains (orange) and shared across all five strains (green). These analyses were performed with the subset of chimeric transcripts with only one TE annotated (1,634).

In both the *overlap and AS insertions* and *internal insertions* groups, TE insertions were also mainly located in the 3’UTRs (53.9%: 316/586; 42%: 503/1199, respectively). In the *internal insertions* group there were more insertions in internal exons than in the *overlap and AS* insertions (414 *vs.* 42; χ² test, *p*-value < 0.001). This pattern also holds when we only consider≥120bp insertions (25 *vs.* 143; χ² test, *p*-value < 0.001; Supplemental Table S4C). Figure 3B shows the number of chimeric gene-TE transcripts globally and by insertion group, body part and strain (Supplemental Table S4D) where it can be observed that, overall, the previous patterns hold at the body part level.

### Retrotransposons and DNA transposons contribute to chimeric gene-TE transcripts

We assessed the contribution of TE families to chimeric gene-TE transcripts. We found that many TE families, 90/146 (62%), were detected in chimeric gene-TE transcripts, as has been previously described in head chimeric transcripts (Supplemental Table S5A; Rech et al. 2022; Treiber and Waddell 2020). Although retrotransposons are more abundant than DNA transposons (60% on average in the five genomes analyzed (Rech et al. 2022), the contribution of retrotransposons to the chimeric gene-TE transcripts was not higher than expected (78%: 70/90; χ² test, *p*-value = 0.14; Supplemental Table S5B). The number of families contributing to the *overlap and AS insertions* group and the *internal insertions* group is similar (75 *vs.* 68, respectively, χ² test, *p-*value = 0.745; Supplemental Table S5C), and both groups were enriched for retrotransposons (χ² test, *p*-values < 0.05; Supplemental Table S5C). Half of these families (46: 51%) contribute to chimeric transcripts in all body parts, while 20 families were body part-specific, with 10 being head-specific, 6 gut-specific and 4 ovary-specific (Supplemental Table S5A).

The most common TE families found were *roo* (38.1%) and INE-1 (26%) (Figure 4). Indeed, these two families were over-represented in the chimeric transcripts dataset when compared to their abundance in the genome: *roo* in the five strains (χ² test, *p*-value < 0.0001 for all comparisons) and INE-1 in SLA-001 (χ² test, *p*-value = 0.0098, respectively) (Supplemental Table S5D). The *roo* and INE-1 were also the most common families both in the *overlap and AS insertions* group (17.3 and 29%, respectively) and in the *internal insertions* group (48.2%and 28.4%, respectively; Figure 4). The same pattern was found when we analyzed only those chimeric transcripts with TEs ≥120bp (Supplemental Fig. S3 and Supplemental Table S5E).

**Figure 4.**
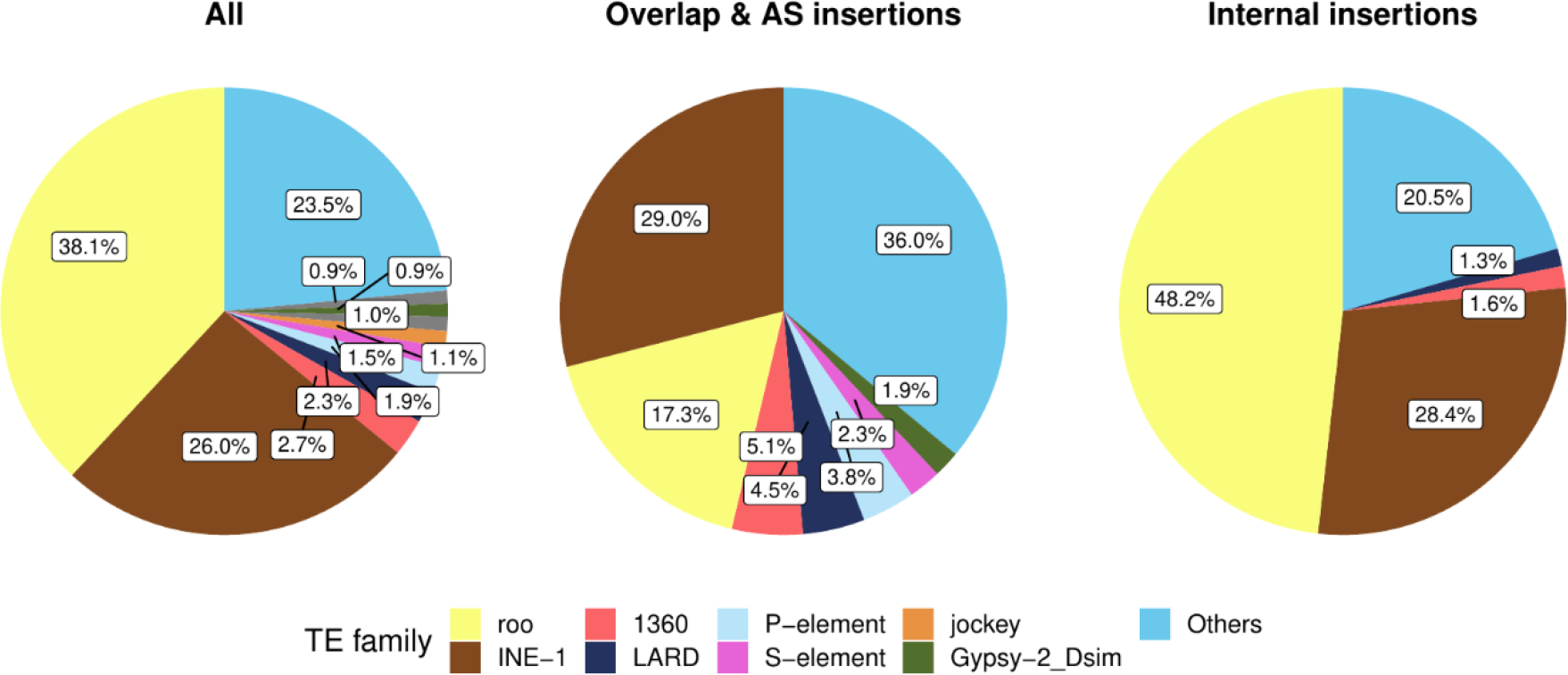
TE families distribution in gene-TE chimeras, globally and by insertion group. Percentage of TE families contributing to gene-TE chimeras in the global dataset (*All*), in the *overlap and AS insertions* group and in the *internal insertions* group. Only TE families found in more than 9 chimeric genes are depicted, otherwise they are grouped in *Others*.

Because *roo* insertions were enriched in all the strains analyzed, we further investigate these TE sequences. We found mainly two types of *roo* insertions: solo LTRs (18 insertions, 16 chimeric transcripts), that most of them belonged to the *overlap and AS insertions* group (16/18), and a short (45bp-192bp) low complexity sequence mapping to the positions 1,052- 1,166 of the canonical *roo* element (see *Methods*). This short *roo* sequence is more common in the *internal insertions* group than in the *overlap and AS insertions* group (2,152 *vs.* 262 insertions, respectively). Note that a recent analysis by Oliveira et al. (2022) also found this same region of the *roo* consensus sequence to be the most abundant in chimeric gene-TE transcripts across four *D. melanogaster* strains. The authors discarded that these short sequences were widespread repeats across the genome as the majority of them (97.45%) were found to have a single blast hit in the genome. We argued that if these low-complexity regions have a *roo* origin, we should find that at least some of them should also have a blast hit with a *roo* insertion. To test this, we used less strict blast parameters compared with Oliveira et al.^43^ and found that 51 of the low complexity regions have a *roo* element insertion as the second best hit and 168 have a *roo* insertion in the top 5 hits, suggesting that indeed some of these sequences have a clear *roo* origin (Supplemental Table S5F). Furthermore, we also tested whether this low complexity region was present in the *roo* consensus sequence from a closely related species, *D. simulans*, and found that this was the case strongly suggesting that this low complexity sequence is an integral part of the *roo* element.

We further investigated why this *roo* low complexity region was incorporated into genes. Because TEs can contain *cis*-regulatory DNA motifs, we performed a motif scan of the low complexity sequence from the canonical *roo* element. We found a C2H2 zinc finger factor motif repeated six times in this region (MA0454.1). Note that this specific motif is only found once in the *roo* consensus sequence outside the low complexity region. A scan in the *roo* sequences from the chimeras revealed that 90% (778/866) of the transcripts with the low complexity *roo* sequence contain at least one sequence of this zinc finger motif, with 25% (196/778) containing 3 or more (Supplemental Table S5G). Additionally, we also look for motifs on *roo* solo LTRs (16 chimeric transcripts) and found that 10 out of 16 transcripts presented at least one motif, all related to DNA-binding domains found in transcription factors, including other C2H2 zinc fingers factor motifs (Supplemental Fig. S4A).

Finally, we also performed a motif scan on INE-1 elements (600 transcripts), the second most common family, and on the 20 body part-specific TE families (29 transcripts). 471 out of 600 of the analyzed transcripts contained motifs provided by the INE-1 fragment, all of them related to DNA-binding motifs, including the C2H2 zinc fingers factors (Supplemental Fig. S4B). 24 out of 29 transcripts with body-specific TE families contained at least one motif related to 16 different classes, including again C2H2 zinc finger factors (Supplemental Fig. S4). Of the 16 motifs, 62.5% (10/16) were body-part specific, with 6 detected in head-chimeric transcripts, 2 in gut and 2 in ovary (Supplemental Fig. S4C).

### Chimeric gene-TE transcripts contribute a median of ∼43% of the total gene expression

Besides identifying and characterizing chimeric gene-TE transcripts, we quantified the level of expression of both chimeric and non-chimeric transcripts genome-wide. We focused on transcripts with ≥1 TPM in at least one of the samples analyzed (1,909 out of 1,931 chimeric transcripts, corresponding to 819/826 of the genes; see *Methods*, Supplemental Table S6A). We found that chimeric gene-TE transcripts have lower expression levels than non-chimeric transcripts (19,228; Wilcoxon’s test, *p*-value < 0.001, Figure 5A). This is in contrast with previous observations in human pluripotent stem cells that reported no differences in expression between chimeric and non-chimeric transcripts (Babarinde et al. 2021). We dismissed the possibility that the lower expression of chimeric gene-TE transcripts was driven by the *roo* low complexity region identified in 829 of the chimeric transcripts (Wilcoxon’s test, *p*-value < 0.0001; Figure 5A). Lower expression of the chimeric gene-TE transcripts compared to non-chimeric transcripts was also found when we analyzed the *overlap and AS insertions* and *internal insertions* groups separately and at the body part and strain levels (Wilcoxon’s test, *p*-value < 0.001 for all comparisons; Figure 5A and Supplemental Table S6B).

**Figure 5.**
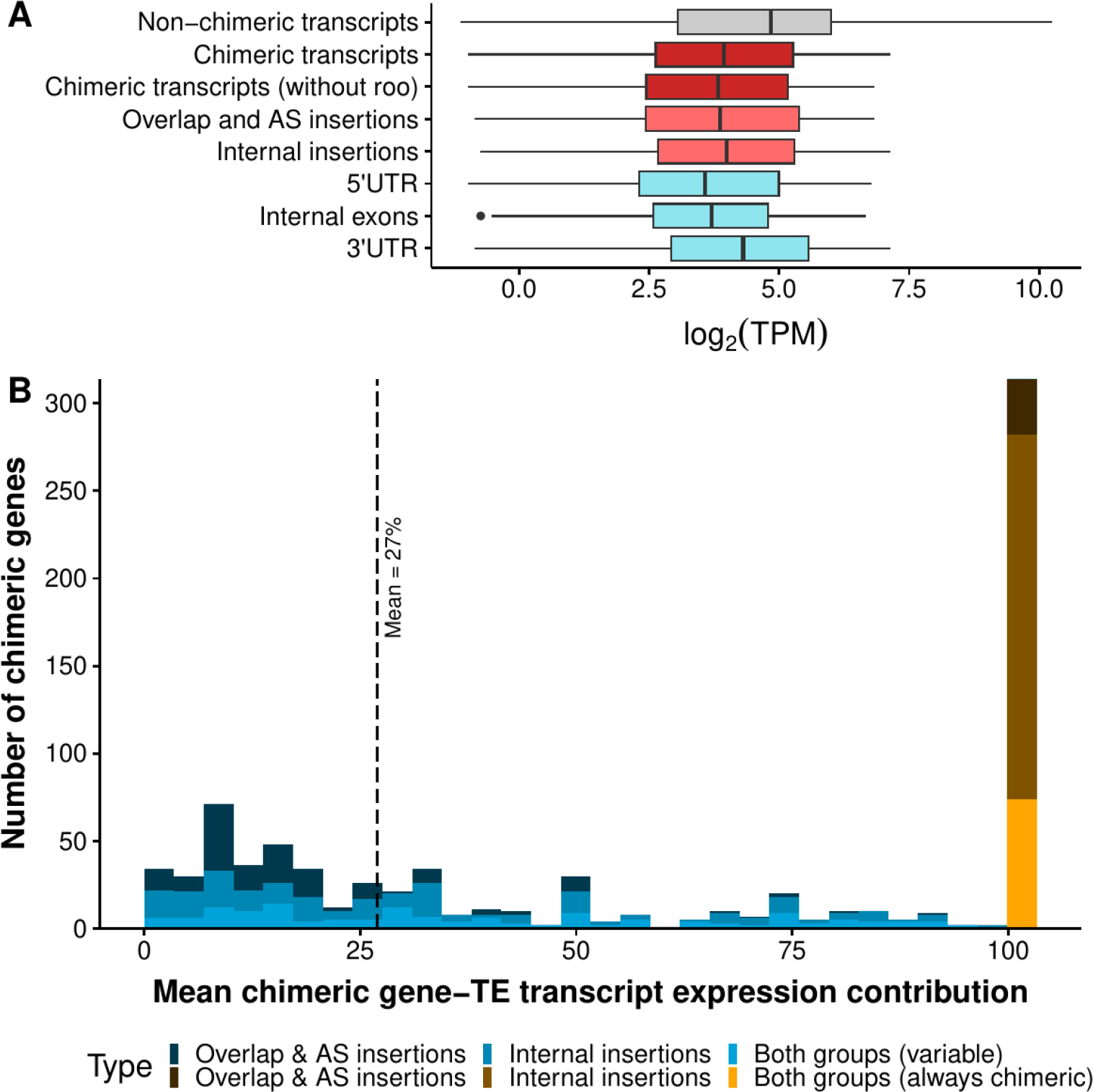
TE insertions within genes affect gene expression. A. Boxplots for the expression levels, measured as the logarithm of TPM: for all non-chimeric transcripts of the genome (19,228, in gray), all chimeric transcripts detected in the present study with TPM ≥1 (1,909, in dark red), chimeric transcripts without the short internal *roo* insertion (1,073, darkred), all chimeric transcripts belonging to the *overlap and AS insertions* group (472, light red) and *internal insertions* group (1,156, light red), and chimeric transcripts divided by position of the insertion (5’UTR: 354, internal exons: 445, 3’UTR: 799, cadet blue). **B.** Histogram showing the expression contribution of chimeric transcripts to the total gene expression. Blue bars represent the contribution of variable chimeric genes (505 genes), ranging from ∼0.1% to >90% (median: 27%) and the orange/brown bar represents the genes that always produced chimeric transcripts in all the genomes and body parts where expression was detected (314 genes).

We further tested whether TEs inserted in different gene locations differed in their levels of expression compared with non-chimeric TE transcripts. We found that chimeric transcripts had significantly lower expression than non-chimeric transcripts regardless of the insertion position (Wilcoxon’s test *p-*value < 0.001 for all comparisons; Figure 5A). Furthermore, insertions in the 3’UTR appeared to be more tolerated than those in 5’UTR and internal exons, as their expression level was higher (Wilcoxon’s test, *p*-value < 0.001 for both comparisons; Figure 5A). Our results are consistent with those reported by Faulkner et al.^22^ who also found that 3’UTR insertions reduced gene expression compared to non-chimeric transcripts and contrast the findings of Babarinde et al. (2021) who found that 3’UTR chimeric transcripts have significantly higher levels of expression compared to non-chimeric transcripts or with insertions in 5’UTR and internal exons.

If we focus on the chimeric genes, 38% of them (314 genes) only expressed the chimeric gene-TE transcript (in all the genomes and body parts where expression of the gene was detected). Most of these genes (76%) contain short TE insertions and accordingly, most of them belong to the *internal insertions* group (66%). For the other 62% (505) of the genes, we calculated the average contribution of the chimeric gene-TE transcript to the total gene expression across samples. While some transcripts contributed on average only ∼0.1% of the total gene expression, others accounted for >90% (mean = 27%) (Figure 5B). The mean contribution to gene expression of the *internal insertions* group was higher than that of the *overlap and AS insertions* group, when considering all the insertions (28.5% *vs.* 15.7%, respectively; Wilcoxon’s test, *p*-value < 0.001), and when analyzing only those transcripts with ≥120bp insertions (28.5 *vs.* 15.7%, respectively; Wilcoxon’s test, *p*-value > 0.0001). Considering only the transcripts that do not contain the *roo* low complexity sequence, the mean contribution to gene expression of the *internal insertions* group was still 25.3%. Overall, taking all chimeric genes into account (819), the average chimeric gene-TE transcripts’ expression contribution to the total gene expression was 43%.

Finally, we evaluated whether there are differences between the expression levels of body part-specific and body part-shared chimeric transcripts. The breadth of expression, measured as the number of tissues in which a gene is expressed, is significantly and positively correlated with the level of expression in *Drosophila* (Larracuente et al. 2008) and humans (Park and Choi 2010). Consistent with this, we found that body part-shared chimeric transcripts have significantly higher expression levels than chimeric transcripts expressed in only one body part (Wilcoxon’s test, *p*-value < 0.001; Supplemental Table S6C), when considering the whole dataset and for chimeric transcripts with insertions ≥120bp (Wilcoxon’s test, *p*-value < 0.001; Supplemental Table S6C). Since we observed that the head was expressing more chimeric transcripts (Figure 2A), we next assessed if head-specific chimeric transcripts were also expressed at higher levels. However, we observed that the median expression of head-specific chimeric transcripts was similar to those specific to gut (medianhead= 9.7 TPM [*n* = 620], mediangut= 10 TPM [*n* = 427]; Wilcoxon’s test, *p*-value = 0.054), but lower than ovary-specific chimeric transcripts (medianovary= 12.6 TPM [*n* = 359]; Wilcoxon’s test, *p*-value > 0.0001). However, this is similar to the expression level of transcripts in these body parts (median of gene expression in ovary>gut>head: 21.2>15.10>12.47).

Interestingly, chimeric transcripts found in all five strains also have significantly higher expression levels than strain-specific chimeric transcripts (Wilcoxon’s test, *p*-value < 0.001; Supplemental Table S6D).

### 17.1% of the TEs within chimeric gene-TE transcripts could also be affecting gene expression via epigenetic changes

We tested whether TEs that produce chimeric transcripts could also be affecting gene expression by affecting the epigenetic marks. We used ChIP-seq experiments previously performed in our lab (Coronado-Zamora et al. 2023) for the three body parts, in each of the five strains analyzed, for two histone marks: the silencing mark H3K9me3 (Choi and Lee 2020; Yin et al. 2011) and the activating mark H3K27ac, related to active promoters and enhancers (Buecker and Wysocka 2012; Koenecke et al. 2016). We tested whether strains expressing the gene-TE chimeric transcript differed in the gene body epigenetic marks compared with strains that do not express the chimeric transcript (430 genes). For the majority of these genes (266), we did not observe consistent epigenetic patterns across samples expressing or not expressing the chimeric transcripts, and these genes were not further analyzed. Additionally, 42 genes did not harbor any epigenetic marks while 93 genes contained the same epigenetics mark(s) (H3K27ac, H3K9me3, or both marks) in strains with and without that particular chimeric transcript (Supplemental Table S7A). Overall, for 17.1% (72/420) of the genes, we observed a consistent change in the epigenetic status associated with the presence of the gene-TE chimera. This percentage is similar for the *overlap and AS* group and the internal insertion group (16.9% and 18.4%, respectively). The majority of TEs showing consistent changes in their epigenetic status were associated with gene down-regulation (68%; Table 1) and this percentage increases when considering chimeric transcripts with insertions in 5’UTR: 88% (16/18) genes are associated with down regulation (Supplemental Table S7B).

**Table 1.** Expression fold-change associated with epigenetic status of strains expressing or not expressing the chimeric transcript. Highlighted in bold, genes showing the expected change in expression according to the gained/lost histone mark.

| Fold change | Non-chimeric gene | No mark | No mark | No mark | H3K27ac | H3K9me3 and H3K27ac | H3K9me3 and H3K27ac | H3K27ac | H3K27ac | H3K9me3 | H3K9me3 and H3K27ac |
| --- | --- | --- | --- | --- | --- | --- | --- | --- | --- | --- | --- |
|  | Chimeric gene | H3K27ac | H3K9me3 | H3K27ac and H3K9me3 | No mark | H3K27ac | H3K9me3 | H3K9me3 | H3K9me3 and H3K27ac | H3K9me3 and H3K27ac | No mark |
| FC > 1 |  | 4 | 1 | 6 | 1 | 3 | 0 | 2 | 4 | 1 | 2 |
| FC < 1 |  | 4 | 2 | 10 | 10 | 1 | 2 | 1 | 14 | 1 | 5 |

### Gene-TE chimeric transcripts are enriched for DNA-binding molecular functions involved in regulation and development

To get insight into the biological processes and molecular functions in which the gene-TE chimeric transcripts are involved, we performed a gene ontology (GO) clustering analysis (Huang et al. 2009). We analyzed the chimeric genes detected in each body part separately, using as a background the total genes assembled in the corresponding body part. We found that chimeric genes are enriched in general cell functions, such as regulation, and development (Figure 6A and Supplemental Table S8A). Some functions are particular to a body part, *e.g.*, *transcription from RNA polymerase II promoter* in the head, *regulation of RNA biosynthetic process* in the gut, and *morphogenesis and cell communication and signaling* in the ovary. Note that the *overlap and AS insertions* group is enriched for biological processes that are body-part specific: *communication signaling and stimulus* in the head, *cellular amide and organitrogen compound metabolic processes* in the gut *and cell projection, assembly and organization* in the ovary (Figure 6A and Supplemental Table S8C).

**Figure 6.**
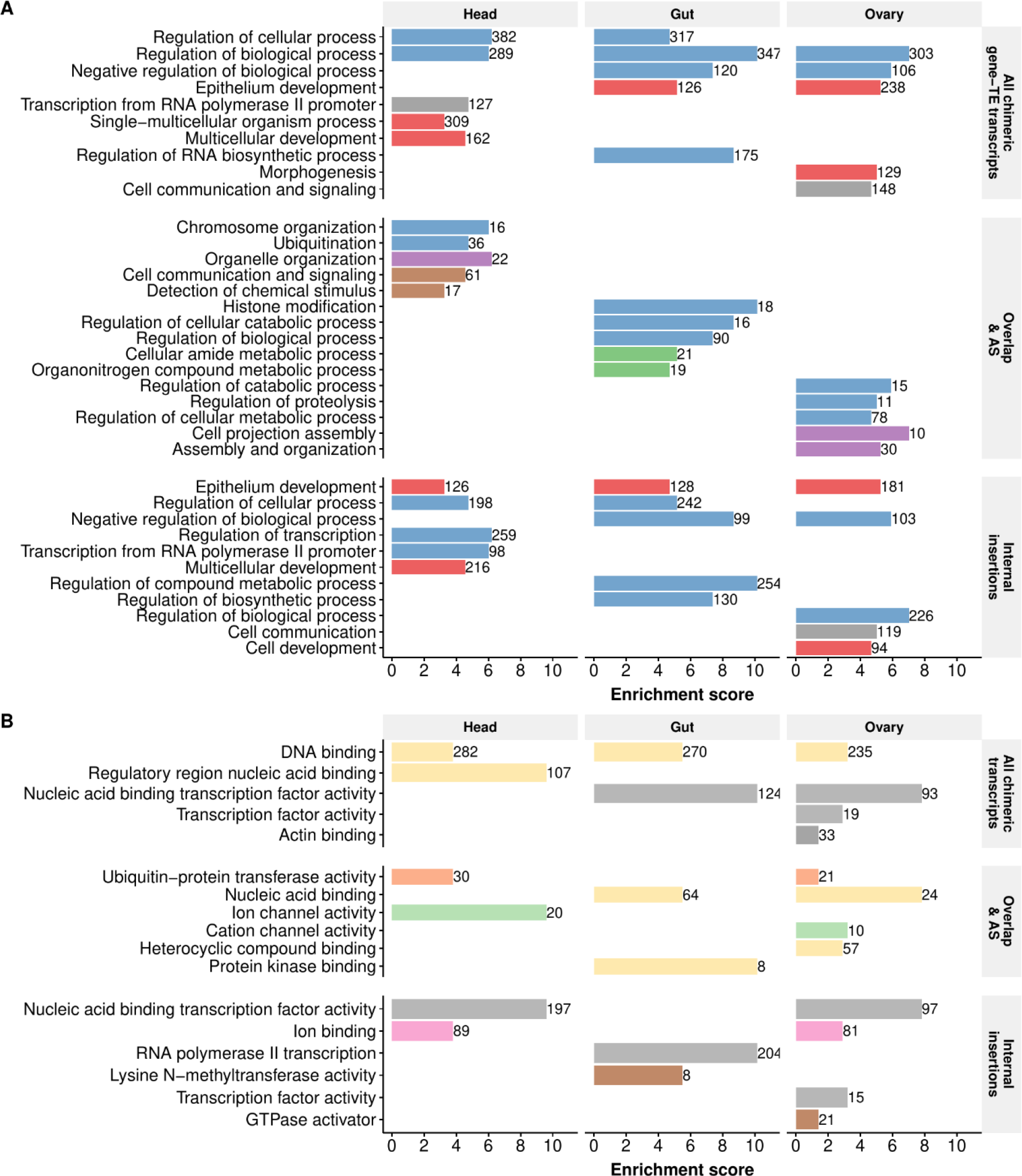
Biological processes and molecular functions of chimeric gene-TE transcripts. A. Biological processes clustering. **B.** Molecular functions clustering. The lengthof the bars represents the cluster enrichment score. The number in the bars represents the number of genes in each cluster. Names of the annotation clusters are manually processed based on the cluster’s GO terms. Colors represent similar annotation clusters. Detailed GO terms of each cluster are given in Supplemental Table S8.

Finally, regarding the molecular function, chimeric genes are enriched for *DNA binding* processes across body parts (Figure 6B and Supplemental Table S8B), while in head they are also enriched for *regulatory region nucleic acid binding* and in ovary for *transcription factor activity* and *actin binding*.

### Both DNA transposons and retrotransposons add functional protein domains

We next assessed whether TE sequences annotated in internal exons provided functional protein domains. We first confirmed, using the Coding Potential Assessment Tool (CPAT; Wang et al. 2013) software, that the majority of chimeric protein-coding gene-TE transcripts that have a TE annotated have coding potential (94.3%: 1,687/1,789; Supplemental Table S9A). Using PFAM (Mistry et al. 2021), we identified a total of 23 PFAM domains in 33 different chimeric transcripts from 24 genes (Table 2 and Supplemental Table S9B and S9C). These 23 domains were identified in 16 TE families, with 9 TE families providing more than one domain. The size of these domains ranged from 9bp to 610bp (median of 72bp; Supplemental Table S9B). Note that 9 of these 24 chimeric genes have been previously described in the literature (Table 2). We observed similar numbers of transcripts belonging to the *Overlap and AS insertions* group and *Internal insertions* group (12 and 16, respectively, with 5 belonging to both). Finally, we found chimeric transcripts adding domains in the three body parts analyzed (Table 2), with an even distribution across them (head: 15, ovary: 13, gut: 12).

**Table 2.**
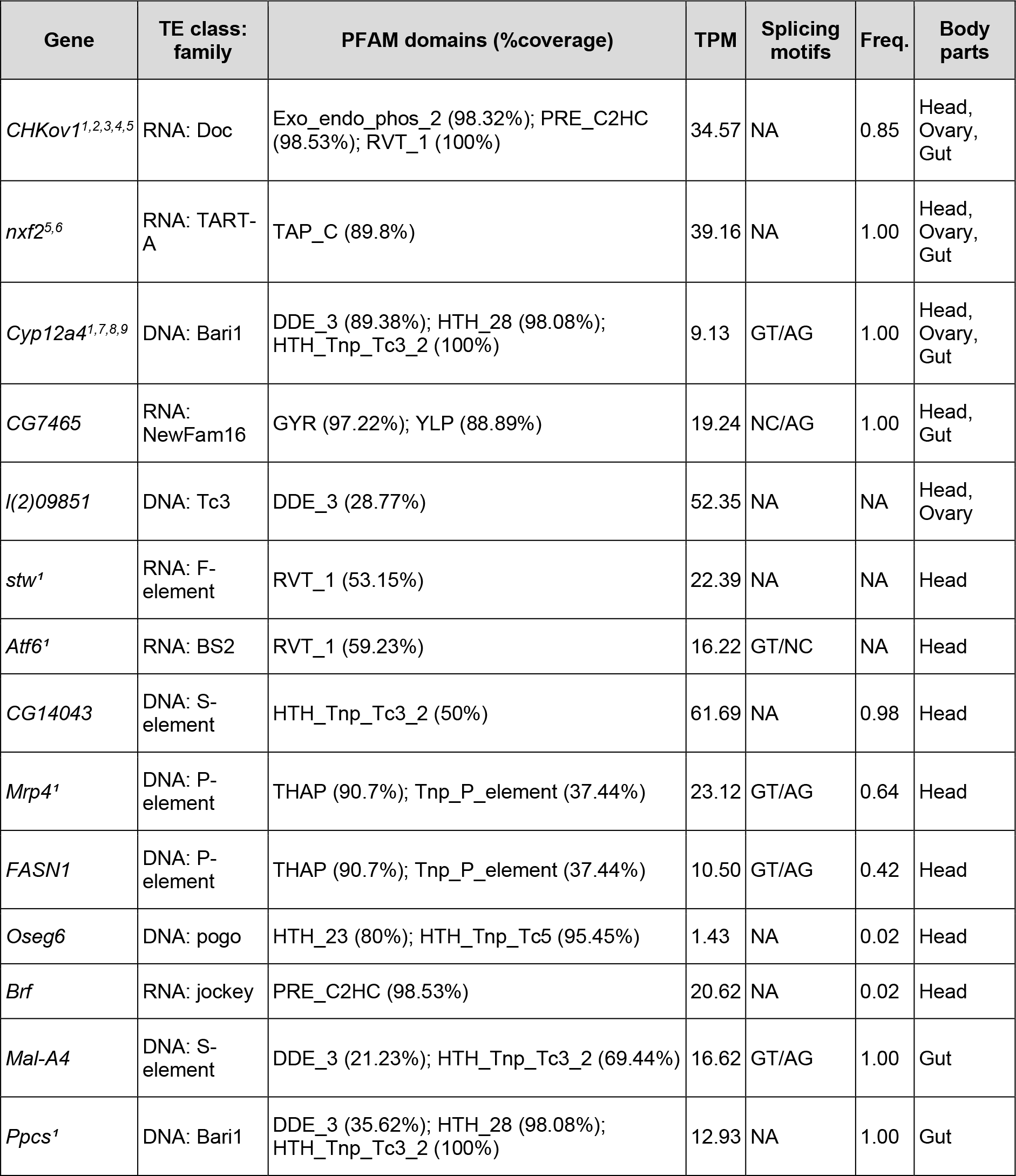
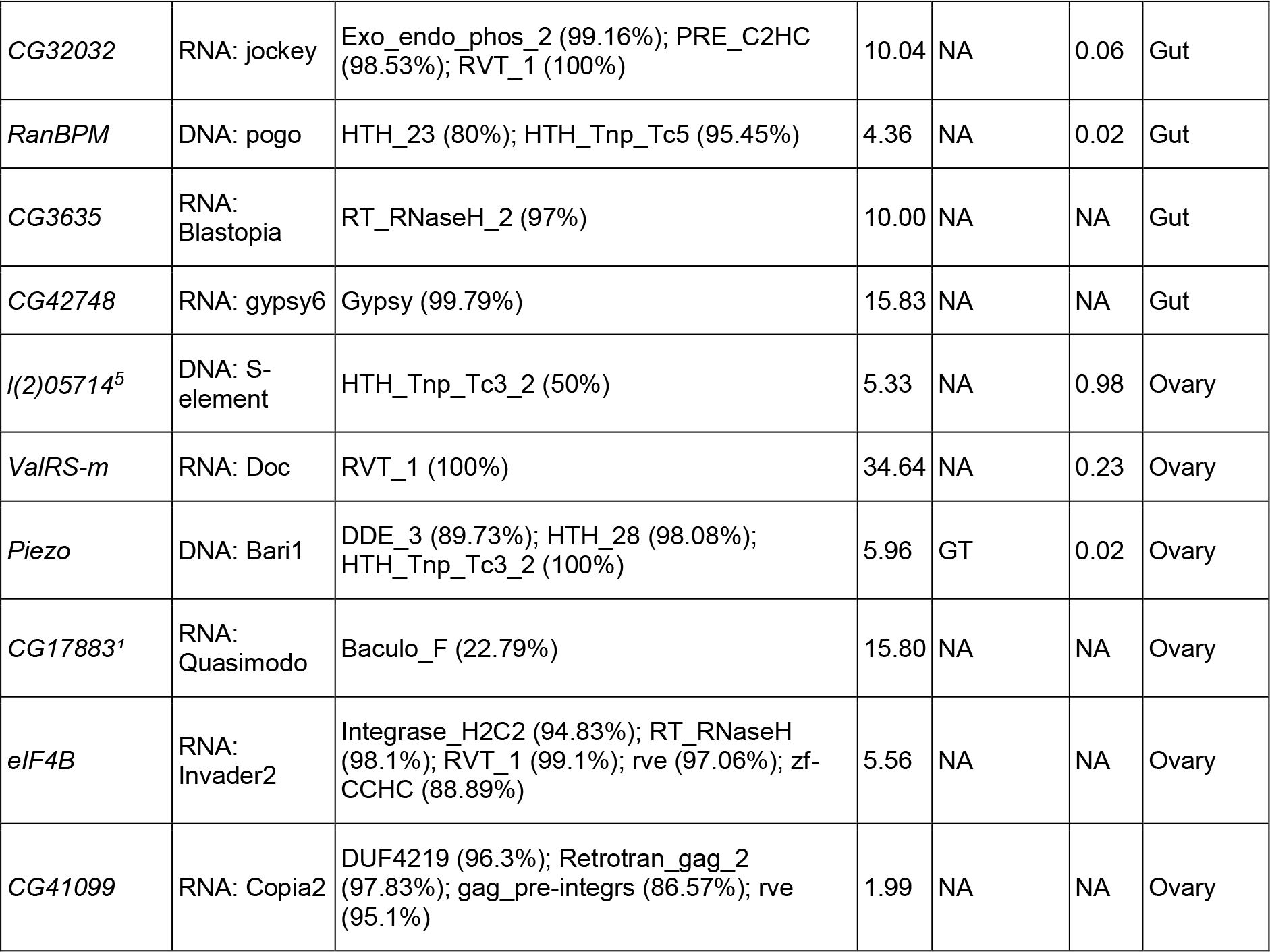
Description of the 24 chimeric genes containing a TE providing a protein domain. Superscript numbers in the *Gene* column represent literature describing these chimeric genes: [^1^] Treiber and Waddell 2020, [^2^] Lipatov et al. 2005, [^3^] Magwire et al. 2011, [^4^] Aminetzach et al. 2005, [^5^] Oliveira et al. 2022, [^6^] Ellison et al. 2020 [^7^] Bogwitz et al. 2005, [^8^] Guio et al. 2018 [^9^] Marsano et al. 2005. TPM is the gene-TE chimera expression level and it is the average across body parts and strains. NA in the *splicing motifs* column represents cases in which there are not splicing signals because the TE was found inside an exon (*internal insertion* group) while NC stands for non-canonical splicing motif. TE frequency (*Freq.*) was retrieved from Rech et al. (2022).

Both DNA transposons and retrotransposons provide domains (13/24 and 11/24, respectively) and most TEs provided a nearly-full domain (22/24, ≥50% coverage), including 6 TEs adding a full-size domain (Table 2). Chimeric genes were related, among others, to transporter functions (11/24), enzyme functions (9/24) and response to stimulus (5/24) (Supplemental Table S9C). All these chimeric genes have evidence of expression, ranging from 1.43 to 61.69 TPM (Table 2, median = 18.16 TPM). We did not observe statistical differences between the level of expression of transcripts with complete domains compared to partially/uncompleted domains (Wilcoxon’s test, *p*-value = 0.535). The majority of TEs for which the population TE frequency has been reported (Rech et al. 2022), are fixed or present at high frequencies (8/14 TEs; Table 2).

We assessed if the domains detected in the TE fragment of the gene-TE chimera were also found in the consensus sequence of the TE family. Because most TE families were providing more than one domain, in total we analyzed 47 unique domains. We were able to find the domain sequence for 44 unique domains from 21 TE consensus sequences (Supplemental Table S9D). Note that for seven of these domains (from seven TEs), we had to lower PFAM detection thresholds to detect them (see *Methods*). The three domains that were not identified in the consensus sequences, were not detected in the chimeric fragments as full domain sequences.

We performed a PFAM domain enrichment analysis considering domains annotated with nearly-full domains and in transcripts expressed with a minimum of 1 TPM using dcGO (Fang and Gough 2013). Overall, seven domains were enriched for the molecular function *catalytic activity, acting on RNA* (4 domains, FDR = 3.44×10^-4^), nucleic acid binding (6 domains, FDR= 3.44×10^-4^), DNA polymerase activity (3 domains, FDR = 6.81×10^-5^) and RNA-directed DNA polymerase activity (3 domains, FDR = 4.66×10^-7^) (Table 3). Five out of seven enriched domains were found in retrotransposon insertions.

**Table 3.** PFAM domain enrichment analysis. dcGO enrichment results using ’Gene Ontology (GO)’ with FDR < 0.01.

| GO term | Z-score | FDR | Annotated domains |
| --- | --- | --- | --- |
| Catalytic activity, acting on RNA | 5.99 | 3.44×10<sup>-4</sup> | PF00078 (RVT_1); PF00098 (zf-CCHC); PF00665 (rve); PF13976 (gag_pre-integrs) |
| Nucleic acid binding | 4.62 | 3.44×10<sup>-4</sup> | PF00098 (zf-CCHC); PF00665 (rve); PF03221 (HTH_Tnp_Tc5); PF13976 (gag_pre-integrs); PF03943 (TAP_C); PF05485 (THAP) |
| DNA polymerase activity | 11.40 | 6.81×10<sup>-5</sup> | PF00078 (RVT_1); PF00665 (rve); PF13976 (gag_pre-integrs) |
| RNA-directed DNA polymerase activity | 27.54 | 4.66×10<sup>-7</sup> | PF00078 (RVT_1); PF00665 (rve); PF13976 (gag_pre-integrs) |

## DISCUSSION

TEs contribute to genome innovation by expanding gene regulation, both of individual genes and of gene regulatory networks, enriching transcript diversity, and providing protein domains (*e.g.*, reviewed in Chuong et al. 2017 and Modzelewski et al. 2022). While the role of TEs as providers of regulatory sequences has been extensively studied, their contribution to transcriptome diversification and protein domain evolution has been less characterized. In this work, we have identified and characterized chimeric gene-TE transcripts across three body parts and five natural *D. melanogaster* strains, and we have quantified their contribution to total gene expression and to protein domains. While previous studies were hindered by the incomplete annotation of TEs in the genome studied (Lipatov et al. 2005; Treiber and Waddell 2020), in this work, we took advantage of the availability of high-quality genome assemblies and genome annotations for five natural strains to carry out an in-depth analysis of gene-TE chimeric transcripts (Rech et al. 2022). We found that TEs contribute ∼9% to the global transcriptome and ∼19% to the body part-specific transcriptome (Figure 1). Contrary to other studies that mostly focus on a single type of chimeric gene-TE transcript, we investigated a comprehensive dataset of chimeras. Indeed, we found that besides insertions affecting the transcription start site, transcript termination, and adding spliced sites (*overlap and alternative splicing insertions*), we also identified a substantial number of TE sequences that were completely embedded within exons (*internal insertions;* Figure 1D). These two types of chimeric gene-TE transcripts shared many properties, *e.g.,* they were enriched for body part-specific transcripts (Supplemental Fig. S2B), and they showed lower expression levels than non-chimeric transcripts (Figure 5A), suggesting that they both should be taken into account when analyzing the contribution of TEs to gene novelty. The *internal insertions* group contributed more to total gene expression (Figure 5B), however, we dismissed the possibility that this increased expression was due to shorter TE insertions, which are more likely to be enriched for false annotations compared with longer insertions (Rech et al. 2022). We found, both based on size and frequency, that the *internal insertions* group is likely to be enriched for older insertions. As such, a higher level of expression of these likely older TEs is consistent with previous observations in tetrapods suggesting that over time gene-TE chimeric transcripts often become the primary or sole transcript for a gene (Cosby et al. 2021). Overall, and taking only into account those gene-TE chimeric transcripts with evidence of expression, we found 102 (5.9%) insertions disrupting the coding capacity, 435 (25.2%) affecting the coding capacity, 316 (18.3%) and 717 (41.6%) affecting the 5’ and the 3’ end of the gene, respectively, while 155 (9%) affected multiple transcript positions (Supplemental Table S6A).

Our finding that TEs contribute to the expansion of the head transcriptome supports the results of Treiber and Waddell (2020) and suggests that ∼6% of genes produce chimeric transcripts in the head due to exonization of a TE insertion. However, because we also analyzed gut and ovary, we further show that TEs can significantly contribute to the expansion of other body parts’ transcriptomes as well (Figure 2). The observation that there are more chimeric transcripts in the head is consistent with a higher transcriptional complexity in the *Drosophila* nervous system tissues (Brown et al. 2014). The fact that chimeric gene-TE transcripts tend to be tissue-specific could be especially relevant for adaptive evolution as tissue-specific genes can free the host from pleiotropic constraints and allow the exploration of new gene functions (Park and Choi 2010; Rogers and Hartl 2012; Salvador-Martínez et al. 2018).

Finally, we identified a total of 23 TE protein domains co-opted by 24 genes (Table 2 and Supplemental Table S9). Nine of these genes have been previously described as chimeric based on high-throughput screenings or individual gene studies, with some of them, *e.g. CHKov1* and *nxf2,* having functional effects (Aminetzach et al. 2005; Ellison et al. 2020; Magwire et al. 2011) (Table 2). The majority of the domains were present in the TE consensus sequences (Supplemental Table S9D). Furthermore, the 23 domains identified were enriched for *nucleic acid binding*, *catalytic activity*, and *DNA polymerase activity* molecular functions (Table 3). Although there is evidence for DNA binding domains being recruited to generate new genes, previous data comes from a comparative genomic approach across tetrapod genomes that focused on DNA transposons as a source of new protein domains (Cosby et al. 2021). The available data for the genome-wide contribution of retrotransposons to protein domains so far is restricted to endogenous retroviruses in mammals (Ueda et al. 2020). In our dataset, which includes both DNA transposons and retrotransposons, the enrichment for DNA binding domains and for catalytic activity is indeed driven by the retrotransposon insertions (Table 2). Although most of the TEs providing protein domains identified in this work for the first time were present at low population frequencies, three were fixed and two were present at high population frequencies and are thus good candidates for follow-up functional analysis (Table 2 and 3).

Although we have detected more chimeric transcripts than any prior *D. melanogaster* study to date, our estimate of the potential contribution of TEs to the diversification of the transcriptome is likely to be an underestimate. First, and as expected, we found that the contribution of TEs to the transcriptome is body part-specific (72.8%, Supplemental Fig. S2B; Conley et al. 2008; Faulkner et al. 2009) and strain-specific (68%, Supplemental Fig. S2A). Thus analyzing other body parts and increasing the number of genomes analyzed will likely identify more chimeric gene-TE transcripts. And second, although our estimate is based on the highly accurate annotations of TE insertions performed using the REPET pipeline (Rech et al. 2022), highly diverged and fragmented TE insertions are difficult to be accurately annotated by any pipeline and as such might go undetected (Gotea and Makałowski 2006; Rodriguez and Makałowski 2022). Still, the combination of an accurate annotation of chimeric gene-TE transcripts, with expression data across tissues, and the investigation of protein domain acquisition carried out in this work, not only significantly advances our knowledge on the role of TEs in gene expression and protein novelty, but also provides a rich resource for follow-up analysis of gene-TE chimeras.

## METHODS

### Fly stocks

Five *D. melanogaster* strains obtained from the European Drosophila Population Genomics Consortium (DrosEU), were selected according to their different geographical origins: AKA- 017 (Akaa, Finland), JUT-011 (Jutland, Denmark), MUN-016 (Munich, Germany), SLA-001 (Slankamen, Serbia) and TOM-007 (Tomelloso, Spain). Further information on collection dates and localities can be found in Rech et al. (2022).

### RNA-seq and ChIP-seq data for three body parts

RNA-seq and ChIP-seq data for the five strains were obtained from Coronado-Zamora et al. (2023). A full description of the protocols used to generate the data can be found in Coronado- Zamora et al. (2023). Briefly, head, gut and ovary body parts of each strain were dissected at the same time. Three replicates of 30 4-6 old-day females each were processed per body part and strain. RNA-seq library preparation was performed using the TruSeq Stranded mRNA Sample Prep kit from Illumina, and sequenced using Illumina 125bp paired-end reads (26.4M- 68.8M reads; Supplemental Table S1). For ChIP-seq, libraries were performed using TruSeq ChIP Library Preparation Kit. Sequencing was carried out in an Illumina HiSeq 2500 platform, generating 50bp single-end reads (22.2M-59.1M reads; Supplemental Table S1). RNA-seq and ChIP-seq raw data are available in the NCBI Sequence Read Archive (SRA) database under BioProject PRJNA643665. The DrosOmics genome browser (http://gonzalezlab.eu/drosomics) compiles all data used in this work and allows for its visual exploration (Coronado-Zamora et al. 2023).

### Transcriptome assembly

#### Reference-guided transcriptome assembly

To perform reference-guided transcriptome assemblies for each body part and strain (15 samples), we followed the protocol described in Pertea et al. (2015) using HISAT2 (v2.2.1, Kim et al. 2019) and StringTie (v2.1.2, Pertea et al. 2015) . We used *D. melanogaster* r6.31 reference gene annotations (Larkin et al. 2021, available at: ftp://ftp.flybase.net/releases/FB2019_06/dmel_r6.31/gtf/dmel-all-r6.31.gtf.gz, last accessed:

October 2020). We first used *extract_splice_sites.py* and *extract_exons.py* python scripts, included in the HISAT2 package, to extract the splice sites and exon information from the gene annotation file. Next, we build the HISAT2 index using *hisat2-build* (argument: *-p 12*) providing the splice sites and exon information obtained in the previous step in the *-ss* and *-exon* arguments, respectively. We performed the mapping of the RNA-seq reads (from the fastq files, previously analyzed with FastQC, Andrews 2010) with HISAT2 (using the command *hisat2 -p 12 --dta -x*). The output sam files were sorted and transformed into bam files using *samtools* (v1.6, Li et al. 2009). Finally, we used StringTie for the assembly of transcripts. We used the optimized parameters for *D. melanogaster* provided in Yang et al. (2018) to perform an accurate transcriptome assembly: *stringtie -c 1.5 -g 51 -f 0.016 -j 2 -a 15 -M 0.95*. Finally, *stringtie --merge* was used to join all the annotation files generated for each body part and strain. We used *gffcompare* (v0.11.2) from the StringTie package to compare the generated assembly with the reference *D. melanogaster* r.6.31 annotation, and the sensitivity and precision at the locus level were 99.7 and 98.5, respectively.

#### De novo transcriptome assembly

A *de novo* transcriptome assembly for each sample was performed using Trinity (v2.15.1, Grabherr et al. 2011) on the fastq files previously processed with *fastp* (v.0.23.2; Chen et al. 2018). Trinity used the following parameters: *--seqType fq --samples_file <TXT file with fastq directory> --CPU 12 --max_memory 78 G --jaccard_clip*. Following Trinity recommended implementation (Haas et al. 2013), we combined all reads across technical replicates, in order to assemble them into a single reference assembly per sample (15 samples: 5 strains × 3 body-parts). The *--jaccard_clip* parameter was used to split falsely fused transcripts derived from gene-dense compact genomes. Next, to keep reliable near full-length transcripts, we used *blastn* (v2.11.0, Camacho et al. 2009) to assign each *de novo* transcript to a known *D. melanogaster* transcript obtained from the *Reference-guided transcriptome assembly*. Next, the script *analyze_blastPlus_topHit_coverage.pl* from Trinity toolkit was used to evaluate the quality of the BLAST results, and we followed a conservative approach that only kept a transcript with a coverage higher than 80% with a known *D. melanogaster* transcript, thus, keeping 150,699 assembled transcripts across all samples. Then, we applied the algorithm proposed by Lima et al. (2017), to detect and remove falsely-fusioned transcripts that were not correctly assembled by Trinity. Briefly, we used *blastn* (Camacho et al. 2009) again to assign each transcript to a known *D. melanogaster* transcript and removed it if it had two or more matches with transcripts from different genes with a coverage of at least 80% and more than 100bp. As expected, most of the fusion transcripts were due to genes overlapping by their UTRs. Finally, we tried to minimize other possible sources of confounding errors by excluding transcripts that were not overlapping a known transcript (tagged by StringTie as *possible polymerase run-on; intron match on the opposite strand [likely a mapping error]; fully contained in a reference intron*; or *intergenic*). The set of final assembled transcripts by Trinity was 131,840.

### Identification and characterization of chimeric gene-TE transcripts

We focused on the set of assembled *de novo* transcripts that passed the previous filters to identify putative chimeric gene-TE transcripts. To annotate TEs in the *de novo* assembled transcripts, we used RepeatMasker (v4.1.2, Smit et al. 2013, with parameters *-norna -nolow - s -cutoff 250 -xsmall -no_is -gff*) with a manually curated TE library (Rech et al. 2022). Note that RepeatMasker states that a cutoff of 250 will guarantee no false positives (Smit et al. 2013). We excluded transcripts for which the entire sequence corresponded to a transposable element, indicative of the autonomous expression of a TE, or TE fragments whose sequence contains simple repeats in more than >80% of their length, as determined by Tandem Repeats Finder (v4.09; Benson 1999), with the exception of *roo*-containing transcripts, that were analyzed separately. To infer the exon-intron boundaries of the transcript, we used *minimap2* (v2.24, Li 2018, with arguments *-ax splice --secondary=no --sam-hit-only -C5 -t4*) to align the transcript to the gene region obtained from the genome of the corresponding strain from which it was assembled. We excluded single-transcript unit transcripts when they matched a multi-exon reference transcript, as it could be indicative of pervasive transcription or non-mature mRNAs. If a single-transcript unit transcript matches a single-exon reference transcript, it is considered as valid. With this process, we obtained the full-length transcript from the genome sequence.

We ran RepeatMasker again (same parameters) on the full-length transcripts to annotate the full TEs and obtain the length of the insertion. We considered that short insertions are those shorter than 120bp (Rech et al. 2022). Finally, we used an *ad-hoc* bash script to define the TE position within the transcript and define the two insertions groups: the *overlap and AS insertions* group and the *internal insertions* group. The *overlap and AS insertions* group have a TE overlapping with the first (5’UTR) or last (3’UTR) exon, or overlap with the exon-intron junction and thus introduce alternative splice sites (see *Splice sites motif scan analysis*). The *internal insertions group* corresponds to TE fragments detected inside exons.

### TE-generating strain-specific chimeras frequency in other strains

To assess the frequency of TE insertions generating strain-specific chimeras, we examined whether the genomes where the chimeric transcript was not identified contained the TE insertion. For that, we used *blastn* (v2.11.0, Camacho et al. 2009) on the gene region, using the TE sequence of the strain-specific chimeric transcript as the query. We applied the following parameters: *-qcov_hsp_perc 80 and -perc_identity 80*. We then categorized the insertions as strain-specific, polymorphic, or fixed.

### Splice sites motif scan analysis

We followed Treiber and Waddell (2020) approach to detect the splice acceptors and splice donor sites in the *alternative splice (AS) insertions* subgroup of chimeric gene-TE transcripts. In brief, we randomly extracted 11-12bp of 500 known donor and acceptor splice sites from the reference *D. melanogaster* r.6.31 genome. Using the MEME tool (v5.4.1, Bailey and Elkan 1994), we screened for the donor and acceptor motifs in these two sequences, using default parameters. The obtained motifs were then searched in the predicted transposon-intron breakpoints position of our transcripts using FIMO (v5.4.1, Grant et al. 2011, with a significant *p*-value threshold of < 0.05).

### *Roo* analyses

#### Identification in the *roo* consensus sequence of the location of the *roo* low complexity region incorporated into gene-TE chimeric transcripts

To determine the location of the *roo* low complexity region in the *roo* consensus sequence, we downloaded the *roo* consensus sequence from FlyBase (version FB2015_02, Larkin et al. 2021, available at https://flybase.org/static_pages/downloads/FB2015_02/transposons/transposon_sequence_set.embl.txt.gz). We extracted the *roo* fragments detected in the chimeric gene-TE transcripts using *bedtools getfasta* (v2.30.0, Quinlan and Hall 2010), and used *blastn* (v2.11.0, Camacho et al. 2009) with parameters -*dust no -soft_masking false -word_size 7 -outfmt 6 - max_target_seqs 1 -evalue 0.05 -gapopen 5 -gapextend 2* to determine the matching position in the consensus sequence.

#### Identification of transcription factor binding sites in *roo* sequences

We retrieved from JASPAR (v2022, Castro-Mondragon et al. 2022) the models for all the transcription factor binding sites (TFBS) motifs of *D. melanogaster* (160 motifs). We used FIMO (v5.4.1, Grant et al. 2011) to scan for TBFS in the repetitive *roo* sequence from the consensus sequence (region: 1052-1166), as well as in the fragments incorporated in the gene-TE chimeras, with a significant threshold of 1×10^-4^.

#### Genome-wide BLAST analysis of *roo* low complexity sequences

We performed a BLAST search with *blastn* (v2.11.0, Camacho et al. 2009) (with parameters: *-dust no -soft_masking false -outfmt 6 -word_size 7 -evalue 0.05 -gapopen 5 -gapextend 2 -qcov_hsp_perc 85 - perc_identity 75*). Next, we used *bedtools intersect* (v2.30.0, Quinlan and Hall 2010) with the gene and transposable elements annotations to see in which positions the matches occur. We analyzed the top 20 matches of each *blastn* search.

#### Identification of *D. simulans roo* consensus sequence

We obtained a superfamily level transposable elements library for *D. simulans* using REPET (Flutre et al. 2011). We used *blastn* (v2.11.0, Camacho et al. 2009) with a minimum coverage and percentage of identity of 80% (*-qcov_hsp_perc 80 -perc_identity 80*) to find the sequence corresponding to the *roo* family. Then, we used again *blastn* (with parameters *-qcov_hsp_perc 80 -perc_identity 80 - dust no -soft_masking false -word_size 7 -max_target_seqs 1 -evalue 0.05 -gapopen 5 - gapextend 2*) to check if the *roo* sequence from *D. simulans* contained the repetitive region present in the *D. melanogaster roo* consensus sequence. The *roo* consensus sequence from *D. simulans* is available in the GitHub repository (https://github.com/GonzalezLab/chimerics-transcripts-dmelanogaster).

#### Retrotransposons and DNA transposons enrichment

We used the percentage of retrotransposons and DNA transposons of the genome of the five strains provided in Rech et al. (2022) and performed a χ² test to compare this percentage to the percentage of retrotransposons and DNA transposons detected in the chimeric gene-TE transcripts dataset.

#### Expression level estimation

The expression level of the whole set of transcripts assembled was computed by Trinity package (v2.15.1, Grabherr et al. 2011), which used *salmon* (Patro et al. 2017) as the abundance estimation method, and applied TPM (transcripts per million) as expression normalization. *Salmon* handles multi-mapped reads and applies an *expectation-maximization* algorithm that attributes read counts to the most likely transcripts. For the expression analyses, we considered transcripts with a minimum expression level of one TPM unless specified. Genes were categorized into three groups: (i) genes that were never detected as producing chimeric isoforms, (ii) genes that always were detected as producing chimeric gene-TE transcripts and (iii) genes producing both chimeric and non-chimeric isoforms. For the later type of genes, we calculated the fraction of the total gene expression that comes from the chimeric transcript to compute the contribution of the chimeric transcripts to the total gene expression.

#### Coding capacity assessment

We assessed whether protein-coding chimeric gene-TE transcripts can produce a protein by using the Coding Potential Assessment Tool (CPAT, v3.0.4) software (Wang et al. 2013) with default parameters. CPAT has been optimized for the prediction of coding and non-coding isoforms in *Drosophila*. Thus, we used the coding probability cutoff at 0.39 (Wang et al. 2013).

#### PFAM scan of domain analysis and enrichment

To scan for PFAM domains (Mistry et al. 2021) in the TEs detected in an internal exon, we extracted the TE sequence from the chimeric transcripts using *bedtools getfasta* (v2.29.2, Quinlan and Hall 2010) translated it to ORF using *getorf* (EMBOSS:6.6.0.0, Rice et al. 2000) and scan it using the script *pfam_scan.pl* (v1.6, Finn et al. 2014) to identify any of the known protein family domains of the Pfam database (version 34). We used dcGO enrichment online tool (Fang and Gough 2013) to perform an enrichment of the PFAM domains detected.

We scanned the consensus TE sequences for the domains present in TE fragments detected in the chimerics transcripts using *pfam_scan.pl* (v1.6, Finn et al. 2014). If the domain was not detected using *pfam_scan.pl* default parameters, we lowered the hmmscan *e*-value sequence and domain cutoffs to 0.05.

#### Chip-seq peak calling

ChIP-seq reads were processed in Coronado-Zamora et al. (2023). Briefly, we used *fastp* (v0.20.1, Chen et al. 2018) to remove adaptors and low-quality sequences. Processed reads were mapped to the corresponding reference genome using the *readAllocate* function (parameter: *chipThres = 500*) of the Perm-seq R package (v0.3.0, Zeng et al. 2015), with *bowtie* (v1.2.2, Langmead et al. 2009) as the aligner and the CSEM program (v2.3, Chung et al. 2011) in order to try to define a single location for multi-mapping reads. In all cases, bowtie was used with default parameters selected by Perm-seq.

Then, we used the ENCODE ChIP-Seq caper pipeline (v2, available at: https://github.com/ENCODE-DCC/chip-seq-pipeline2) in *histone* mode, using *bowtie2* as the aligner, disabling pseudo replicate generation and all related analyses (argument *chip.true_rep_only = TRUE*) and pooling controls (argument *chip.always_use_pooled_ctl = TRUE*). MACS2 peak caller was used with default settings. We used the output narrowPeak files obtained for each replicate of each sample to call the histone peaks. To process the peak data and keep a reliable set of peaks for each sample, we first obtained the summit of every peak and extended it ±100bp. Next, we kept those peaks that overlapped in at least two out of three replicates (following Yang et al. 2014) allowing a maximum gap of 100bp, and merged them in a single file using *bedtools merge* (v2.30.0, Quinlan and Hall 2010). Thus, we obtained for every histone mark of each sample a peak file. We considered that a chimeric gene-TE transcript had a consistent epigenetic status when the same epigenetic status was detected in at least 80% of the samples in which it was detected.

#### GO clustering analysis

The Gene Ontology (GO) clustering analysis in the biological process (BP) and molecular process (MP) category was performed using the DAVID bioinformatics online tool (Huang et al. 2009). Names of the annotation clusters were manually processed based on the cluster’s GO terms. Only clusters with a score >1.3 were considered (Huang et al. 2009).

### Statistical analysis

All statistical analyses were performed in R (v4.1.2) statistical computing environment (R Core Team 2021). Graphics were created using *ggplot2* R package (Wickham 2016).

## DATA ACCESS

The set of chimeric gene-TE transcripts detected in this work are available in GitHub (https://github.com/GonzalezLab/chimerics-transcripts-dmelanogaster). Scripts to perform analyses are available at GitHub (https://github.com/GonzalezLab/chimerics-transcripts-dmelanogaster).

## ACKNOWLEDGMENTS

We thank Carlos Vargas-Chavez and Simón Orozco for providing the *D. simulans* REPET TE library. We thank Simón Orozco and Ewan Harney for comments on the manuscript. This project has received funding from the European Research Council (ERC) under the European Union’s Horizon 2020 research and innovation programme (H2020-ERC-2014-CoG-647900), and from grant PID2020-115874GB-I00 funded by MCIN/AEI/10.13039/501100011033.

## AUTHOR CONTRIBUTIONS

JG conceived the project; JG and MCZ designed the analyses; MCZ performed the analyses; JG and MCZ wrote and revised the manuscript.

