## Supplementary figures and images for "Transposons contribute to the functional diversification of the head, gut, and ovary transcriptomes across *Drosophila* natural strains"

### Supp Fig 1

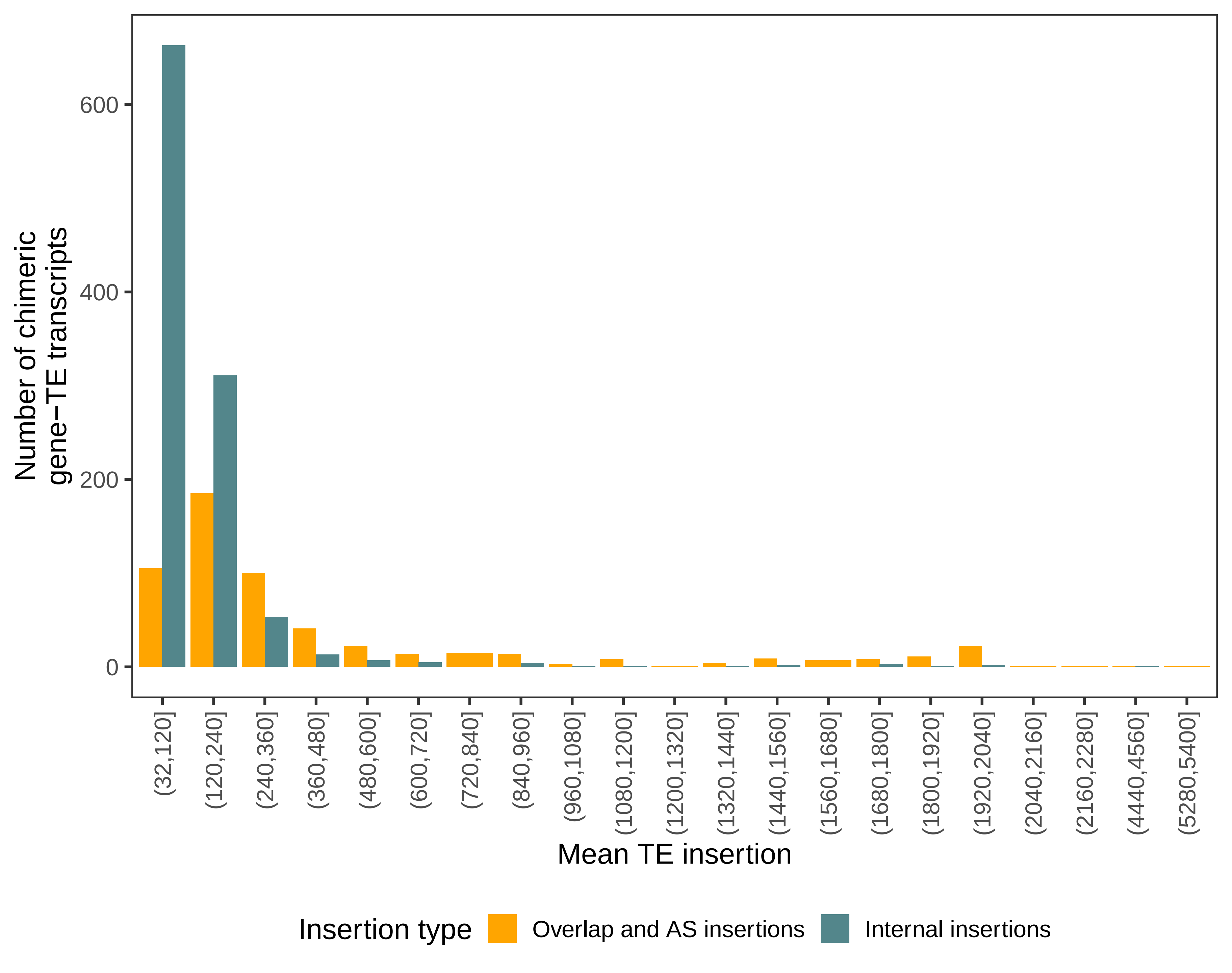

### Supp Fig 2

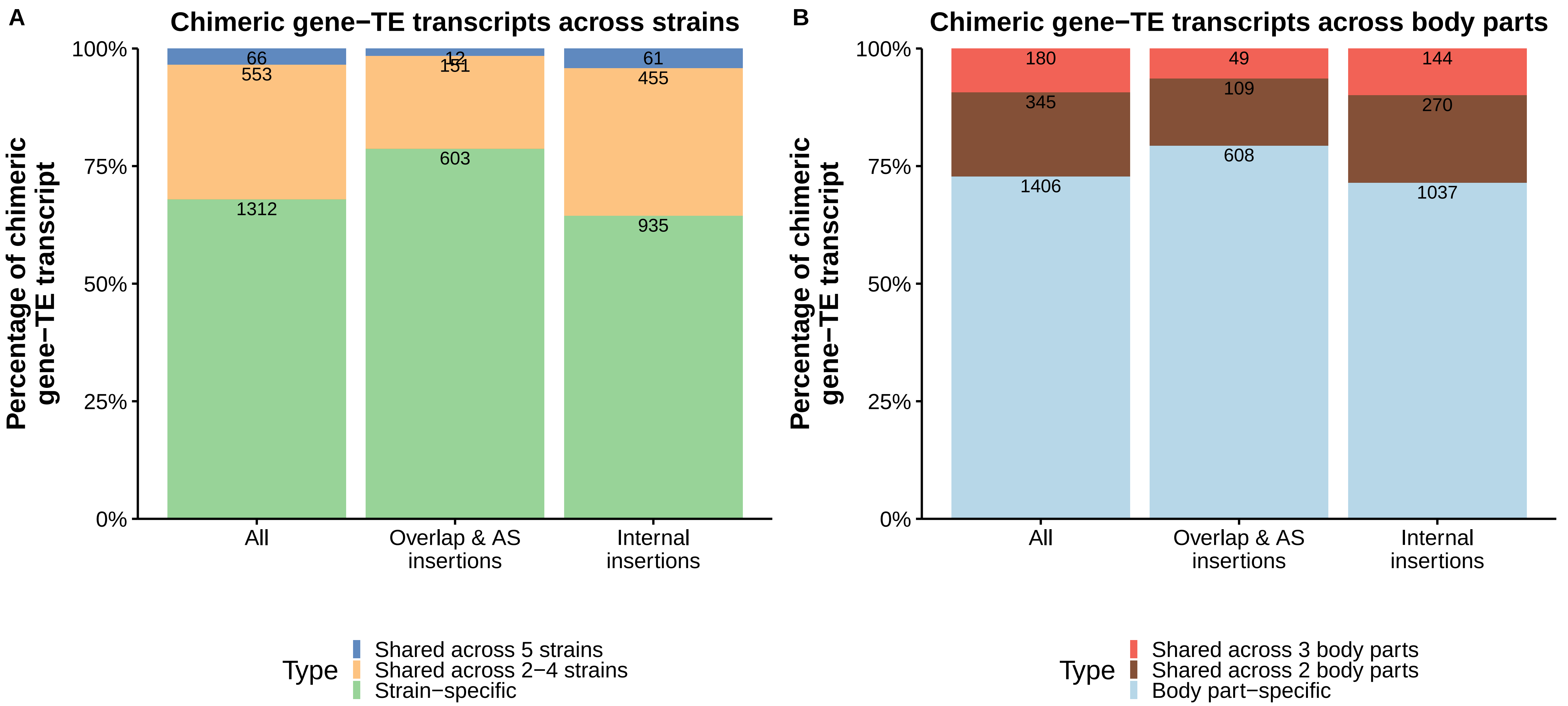

### Supp Fig 3

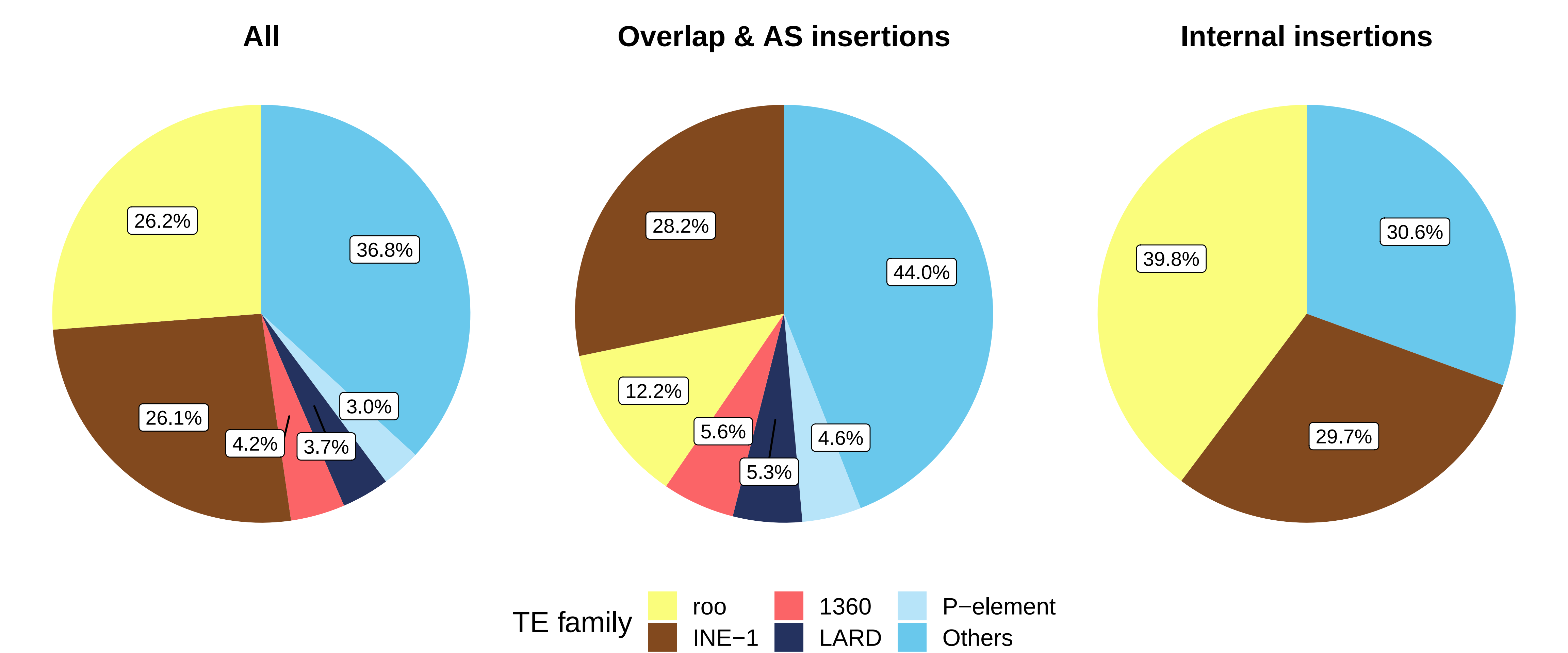

### Supp Fig 4

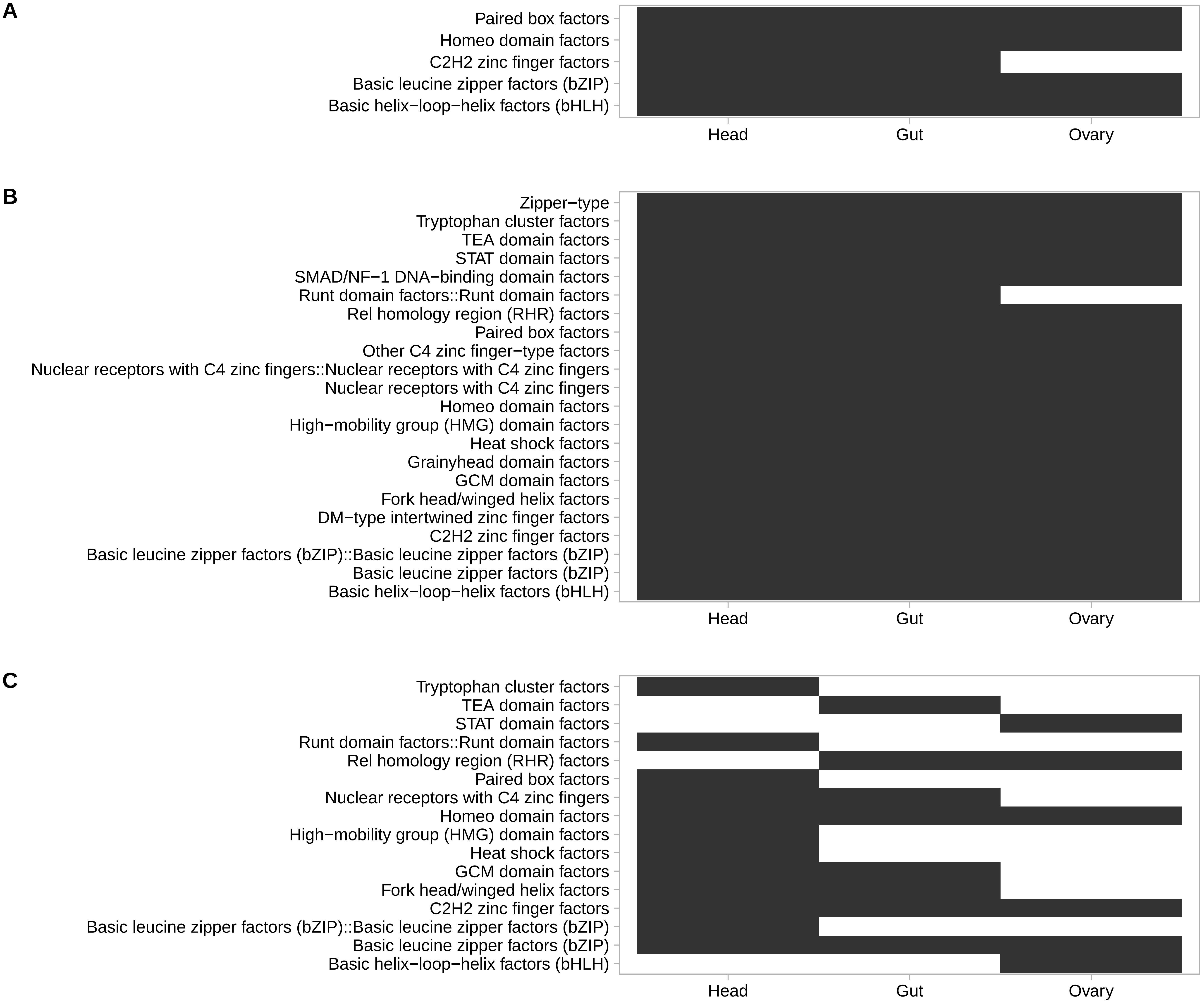
